# SERPINB13 is a prognostic biomarker for LUSC associated with an immune-inflamed tumor phenotype and modulated by immune cells

**DOI:** 10.64898/2026.07.11.737907

**Authors:** Christiane Kuempers, Karina Stein, Doerte Nitschkowski, Tobias Jagomast, Carsten Heidel, Jutta Kirfel, Daniel Droemann, Sabine Bohnet, Michael Schweigert, Martin Reck, Till Olchers, Soenke von Weihe, Ole Ammerpohl, Torsten Goldmann

## Abstract

Non-small cell lung cancer (NSCLC) is the most common form of lung cancer accounting for most cancer-related deaths worldwide. Despite substantial recent advances in targeted therapies and immunotherapy, the prognosis for advanced-stage disease remains comparably poor, which is why the identification of novel molecular biomarkers as well as therapeutic targets influencing tumor development, progression, and metastasis remain important. This study focusses on SERPINB13, a serine-protease inhibitor expressed in selected tissues that is dysregulated in several tumor entities. However, its role in NSCLC still remains largely unclear.

We analyzed *SERPINB13* transcription in a cohort of non-small cell lung cancer (NSCLC) cases including both lung squamous cell carcinoma (LUSC) and lung adenocarcinoma (LUAD) by transcriptome profiling. Epigenetic modifications were assessed via Methylation BeadChips. Additionally, SERPINB13 protein expression was assessed by immunohistochemistry (IHC) in an independent cohort of NSCLC comprising 126 LUSC patients. Correlation analyses were performed to associate SERPINB13 expression with key clinico-pathological parameters, including overall survival and extent of tumor-infiltrating immune cells. To functionally investigate the regulatory influence of peripheral blood mononuclear cells (PBMCs) on *SERPINB13* expression in LUSC tumor cells *in vitro*, we utilized the *SERPINB13*-expressing LUSC cell line LUDLU-1. Here, gene transcription was analyzed by quantitative real-time PCR (RT-qPCR), confirmed by Western blot on the protein level.

Transcriptome analysis revealed a significant upregulation of *SERPINB13* in lung squamous cell carcinoma (LUSC) compared to lung adenocarcinoma (LUAD), highlighting a subtype-specific expression pattern. This differential expression was further associated with a distinct epigenetic DNA methylation signature at the SERPINB13 loci in LUSC, suggesting transcriptional regulation via hypomethylation. IHC analysis demonstrated that high SERPINB13 protein expression is significantly associated with prolonged overall survival in LUSC. Notably, *SERPINB13* expression was enriched in immune-inflamed (“hot”) tumors, characterized by elevated infiltrating lymphocytes and immune activation. Mechanistically, co-culture experiments with PBMCs induced *SERPINB13* expression in a LUSC cell line in a dose- and time-dependent manner in the absence of direct cell contact. This suggests that soluble factors secreted by immune cells might play a key role in regulating *SERPINB13* expression in the tumor microenvironment.

Taken together, SERPINB13 is a novel prognostic indicator in LUSC that is modulated by Immune cells. Further studies are necessary to decipher the crosstalk of Immune cells on the Serpin B13 expressing tumor cells in depth, with regard to a possible interventional strategy. immunomodulatory potential strategies and personalized therapeutic approaches in NSCLC.

## Introduction

Over the past decade, the discovery of various gene alterations as well as immune checkpoints has led to the development of a multitude of novel prognostic factors and therapies in NSCLC. Patients suffering from LUAD have benefited more from the advent of molecular targeted therapies than patients suffering from LUSC, in which progress in the treatment is still limited, resulting in a poorer prognosis ^1^. This underscores the urgent need for novel molecular biomarkers and therapeutic targets especially for this entity. In the context of new therapeutic options, especially immunotherapy, molecules and proteins involved in the dynamic crosstalk between tumor cells and the tumor immune microenvironment (TIME) are gaining increasing attention. Even beyond the context of immunotherapy, the tumor-associated immune infiltrate is known to have a significant impact on the prognosis of NSCLC ^2, 3^.

The inhibitory effect of SERPINs (acronym for serine protease inhibitor) on proteases regulates multiple biological processes, such as inflammation, apoptosis, and cellular stress responses^4^. SERPINB13 (Hurpin, Headpin, PI13) is primarily expressed in squamous epithelia, beinginvolved in processes such as epidermal differentiation, epidermal homeostasis and hair development^5^. Increased levels of SERPINB13 protein have been observed in inflammatory skin diseases such as psoriasis, as well as in (pre)malignancies of the skin like actinic keratosis and squamous cell carcinoma^6^. Among other genes of the SERPIN family *SERPINB13* is located at chromosome 18q21, a region where loss of heterogeneity (LOH) has been observed in malignancies such as squamous cell carcinomas (SCC) of head and neck (HNSCC) and oral cavity (OSCC) ^7, 8^.

A study by Shiiba et al. analyzed mRNA expression of *SERPINs* at this chromosome region in OSCC (n=55) showing a significant decrease in *SERPINB13* expression^8^. De Koning et al. investigated HNSCC and observed partially or totally suppressed SERPINB13 protein in 75% of the samples (n=99), compared with endogenous expression in non-neoplastic epithelial cells which was associated with poor clinical outcomes, including low tumor differentiation, the presence of lymph node metastases, and an unfavorable disease-free survival (DFS) and overall survival (OS). In addition, they found a significant correlation between reduced SERPINB13 protein levels and the presence of LOH at the serpin cluster on chromosome 18q21.3^7^.

These studies suggest SERPINB13 to function as an important protease inhibitor involved in the progression of SCC in general. Its expression and role in NSCLC have been barely studied so far, and, to the best of our knowledge, little is known about protein expression of SERPINB13 in NSCLC.

However, there are studies dealing with SERPINB13 based on *in silico* data. Relli et al. investigated sets of differentially expressed genes in LUAD versus LUSC which were assessed for prognostic impact through meta-analysis of DNA microarray data from 2.437 NSCLC patients through the KMPlot database. An independent analysis of a case series of 994 NSCLC was conducted by next-generation sequencing, together with gene expression profiling from GEO (https://www.ncbi.nlm.nih.gov/geo/). They found SERPINB13 bearing a negative prognostic impact on LUAD, but a positive one in LUSC^9^. A recent study based on machine learning algorithms reports *SERPINB13* to be one of the top genes differentially expressed between LUAD and LUSC^10^. Considering that SERPINB13 participates in the regulation of keratinocyte proliferation and differentiation, it is reasonable to speculate its potential role in LUSC ^5,10^.

Zhan et al. identified *SERPINB13* as one of fifteen genes greatly elevated in LUSC using high-throughput data obtained from The Cancer Genome Atlas (TCGA) database ^11^.

A further study by Zhang et al. focusses on prognostic stratification and therapeutic targeting in LUSC via integrative multi-omic and machine learning approaches, while also considering tumor immune infiltration. The authors constructed a subtype-specific machine learning model called LUSC multi-omics signature (LMS) through a combination of ten machine learning algorithms which demonstrated superior predictive performance than previously published LUSC biomarkers. Patients with lower LMS scores showed longer OS and better responses to immunotherapy compared to patients displaying high LMS scores, while the latter group also had a higher incidence of immune suppression and exclusion (“cold” tumors). *SERPINB13* was identified as one gene showing a significant positive correlation with OS, DFS and PFS of LUSC patients. Notably, a negative correlation between *SERPINB13* expression levels on the RNA-level and the immune score was stated. *In vitro* analyses based on RT-qPCR confirmed that *SERPINB13* was expressed at higher levels in LUSC tissues compared to adjacent non-cancerous tissue. Moreover, the role of SERPINB13 as a potential oncogene was confirmed in NCI-H520 LUSC cells. CCK-8 experiments revealed a notable enhancement in cell viability following *SERPINB13* knockout, and colony formation assays demonstrated a significant increase in colony numbers in cells with reduced *SERPINB13* expression and rate of colony formation was accelerated in *SERPINB13* knockdown cells, suggesting a potential inhibitory role of *SERPINB13* in cell proliferation. Furthermore, migration and invasion assays revealed significantly enhanced capabilities in *SERPINB13* knockdown cells with substantial increase of migrating cells with faster wound healing rate, underscoring a tumor suppressive role of *SERPINB13* in LUSC ^12^.

In summary, *SERPINB13* expression might play a distinct biological role in LUSC pathogenesis, potentially linked to immune modulation and epigenetic regulation. Therefore, we comprehensively analyzed SERPINB13 in two cohorts of NSCLC patients focusing on LUSC, at the epigenetic, RNA, and protein level. We investigated its association with clinic-pathological parameters, especially survival and tumor immune cell infiltration, and its regulation by peripheral blood mononuclear cells (PBMCs) to address the question whether SERPINB13 may function as prognostic biomarker for LUSC and as a dynamic sensor of immune activity in the tumor microenvironment.

## Material and Methods

### Tissues for methylome and transcriptome analysis

We collected tissue samples from NSCLC patients undergoing curative surgical removal of lung cancer. The samples were analyzed together with their matched tumor-free lung tissues. The protocol was approved by the local ethics council at the University of Luebeck (AZ-12-220). Tumor samples were macro-dissected to enrich the tumor content for methylome and transcriptome analyses.

### Methylome analysis of the SERPINB13 gene in NSCLC

Epigenetic modifications of the SERPINB13 gene were analyzed in patients with LUSC (n=18; 14 males, 4 females) and LUAD (n=14; 5 males, 9 females) by Infinium Human Methylation450k BeadChips (Illumina Inc., San Diego, CA, USA) as previously described ^13^. The mean age of the patients at surgery was 65.6 years, 16 were smokers, 14 were ex-smokers, one patient was a never smoker, and smoking status was unknown in one case. Tumor stages were divided according to the 8th Edition of the Union Internationale Contre le Cancer (UICC) staging system as follows: T1 (n=4), T2 (n=18), T3 (n=7), T4 (n=3). Hierarchical clustering of methylation levels of Serpin B13 CpG loci obtained by GenomeStudio software was visualized using the Omics Explorer 3.2 (Qlucore, Lund, Sweden).

### Transcriptome analysis of SERPINB13 in NSCLC

*Transcriptome analysis was performed on a* primary NSCLC collective of 18 NSCLC comprising 10 LUAD and 8 LUSC, respectively, deriving from the cohort used for Methylome analysis. This subcohort consisted of 9 female and 9 male patients. The mean age at diagnosis was 66.06 years, 10 were smokers, 7 were ex-smokers, and one patient was a never-smoker. Tumor stages were divided according to the 8th Edition of the UICC staging system as follows: T1 (n=1), T2 (n=9), T3 (n=6), T4 (n=2).

The relative Serpin B13 gene expression values at the RNA level were quantile-normalized and extracted from the GEO-dataset GSE74706, then analyzed with GraphPad Prism v.8.

### Immunohistochemistry

For immunohistochemistry (IHC), tissue microarrays (TMA) were constructed from formalin-fixed paraffin-embedded (FFPE) tumor containing blocks. For each case, three 1.2 mm diameter cores were obtained from representative tumor regions and transferred into recipient paraffin blocks using a manual tissue arrayer. In some cases, adjacent normal lung tissue was also incorporated into the TMAs. A tumor sample was incorporated in further analysis if at least one core was evaluable.

### SERPINB13 protein expression assessed by IHC

To perform the IHC staining, FFPE tissue sections were deparaffinized, rehydrated and subjected to heat-induced epitope retrieval in Tris buffer (pH 8.4; 4 minutes at 92 °C). Endogenous peroxidase activity was inhibited with 3% hydrogen peroxide for 10 minutes at room temperature. Sections were incubated for 32 minutes at 37°C with a polyclonal rabbit anti-SERPINB13 (SigmaAldrich, HPA057129; dilution 1:250). Detection was performed using a horseradish peroxidase (HRP)-polymer (anti-mouse/rabbit; 20 minutes, room temperature). Sections were then counterstained with Mayer’s hematoxylin for 8 minutes, blued with bluing reagent (Roche Ventana) and finally dehydrated through a graded ethanol series, cleared in xylene and mounted using Pertex.

The immunohistochemically stained samples within the TMAs were microscopically assessed by conventional visual inspection with a light microscope (Olympus BH-2). Staining pattern was cytoplasmic. Staining intensity was categorized into four levels: negative, weak, moderate, and strong. For additional dichotomous analyses, specimens with negative and weak staining were classified as low, and those with moderate or strong staining were classified as high.

### Evaluation of Immune cell infiltration

Cancer samples were classified into those displaying heavy immune cell infiltration (“hot”) and those showing low immune cell infiltration (“cold”), as previously described ^14^. In short, tumor samples with less than 150 lymphocytes per HPF were classified as “cold” while tumor samples with 150 or more lymphocytes per HPF were classified as “hot”.

We further characterized several immune cells subtypes within the tumor immune microenvironment via IHC and stained CD68 (ready-to-use VENTANA Roche anti-CD68 (KP-1), CD3 (ready-to-use VENTANA Roche anti-CD3 (2GV6), CD20 (ready-to-use VENTANA Roche anti-CD20 (L26), CD8 (ready-to-use VENTANA Roche anti-CD8 (SP57, CD4 (ready-to-use VENTANA Roche anti-CD4 (SP35) and PD-L1 (Dako, clone 22C3). IHC was performed by using the Roche Ventana Technology Benchmark Ultra IHC/ISH System (Ventana MedicalSystems, Tucson, AZ, USA).

Macrophages and CD3-, CD20-, CD8- and CD4-positive lymphocytes were counted in three high power fields (HPFs) per core, meaning that up to nine HPFs per case were assessed, as previously described ^15^.

PD-L1 expression on tumor cells was assessed by taking membranous staining into account and indicated as tumor proportion score (percentage of PD-L1 positive tumor cells of all tumor cells; TPS). For the PD-L1 immune score (IS), the proportion of PD-L1-positive immune cells per core was estimated, as described previously ^16^.

### Cohort of IHC analysis

126 patients suffering from LUSC undergoing surgical resection were enrolled in this study as an independent cohort. The median age of the patients (43 female, 83 male) at initial diagnosis was 68 years. Tumors were graded according to the 2015 World Health Organization Classification of Lung Tumors. For determination of tumor state, the 8^th^ and 9^th^ Edition of UICC/TNM staging system was used. From primary tumors, 46 (36.5%), 41 (32.5%), 22 (17.5%) and 16 (12.7%) were classified as pT1, pT2, pT3 and pT4, respectively. In 1 case (0.8%) note concerning T-stage was missing. Archived tissue blocks and slides were collected from 2013 to 2017. All data were anonymized before inclusion in this retrospective study cohort. This study was approved by the Internal Review Board of the University of Luebeck (AZ 16-277).

### Co-culture

LUDLU-1 were seeded on the membrane of 24-well cell culture inserts (0.4 µm pores, TP, PET, Sarstedt, Nümbrecht, Germany) with 0.05 x 10^6^ cells in 200 µl RPMI 1640 (Thermo Fisher Scientific, Waltham, Massachusetts, USA) supplemented with 10 % fetal calf serum (FCS) and 1 % penicillin/streptomycin (PS), further referred to as complete medium. In addition, 500 µl complete medium was transferred in the well under the insert and cells were incubated at 37 °C under 5% CO₂ for 48 hours. Blood of healthy donors was prepared by density gradient centrifugation using Pancoll (PAN Biotech, Aidenbach, Germany) as described elsewhere, and peripheral blood mononuclear cells (PBMC) were harvested and extensively washed with PBS ^17^. When LUDLU-1 reached 80–90% confluence, complete medium was replaced and PBMCs were added in the well under the membrane with a concentration of 0.5 x 10^6^, 0.2 x 10^6^, and 0.05 x 10^6^ cells in 500 µl complete medium, respectively. The co-cultures were incubated for additional 24 and 120 hours with replacing the medium above the membranes every other day.

### Western Blot

LUDLU-1 cells were lysed in 150 µl 1x NuPAGE™ LDS Sample Buffer (Thermo Fisher Scientific, Waltham, Massachusetts, USA) and heated for 10 min at 70°C. Proteins were separated by SDS-PAGE using a 10 % SDS gel. Blotting was performed by using the Power Blotter System with Power Blotter Select Transfer Stacks (Thermo Fisher Scientific). Membranes were blocked with ROTI^®^Block (Roth, Karlsruhe, Germany) for 1 h at room temperature and washed with Tris-buffered saline with 0.1% Tween ® 20 (TBS-T). Membranes were then incubated over night with primary antibodies diluted in ROTI^®^Block: anti-SERPINB13 (1:1000, rabbit polyclonal, #HPA057129, Merck, Darmstadt, Deutschland) and anti-β-Actin (1:5000, mouse monoclonal, # 3700S, Cell Signaling, Danvers, Massachusetts, USA) as a loading control. After 4 additional washing steps with TBS-T, the membranes were incubated with secondary antibodies goat anti-mouse IRDye® 800CW and donkey anti-mouse IRDye® 680RD (#929-70020 and #926-68072, 1:10,000, LI-COR, Lincoln, Nebraska, USA) for 45 min in the dark. Visualization was done with the Odyssey® CLx Imaging System (LI-COR). Quantification was done using the band intensity analysis tool from the software Image Studio Light V5.25 (LI-COR). For that, mean band intensities were measured, and relative intensity was calculated by dividing SERPINB13 values to GAPDH values. Finally, relative values were normalized to the respective controls.

### Immunofluorescence staining for confocal microscopy

LUDLU-1 cells were seeded into 8-well μ-slides (ibidi GmbH, Gräfelfing, Germany) at a density of 300,000 cells per well in complete medium (RPMI-1640 supplemented with 10 % FCS and 1 % PS). Cells were incubated at 37 °C in a humidified atmosphere containing 5 % CO₂ for 24 h. Medium was aspirated, and cells were washed once with phosphate-buffered saline (PBS). Cells were fixed with 4 % paraformaldehyde (BioLegend, San Diego, CA, USA) for 30 min, followed by one wash with PBS. Permeabilization was carried out with 0.25 % Triton X-100 in PBS for 5 min, followed by washing with PBS. Non-specific binding was blocked by incubation with 1 % bovine serum albumin (BSA) in PBS for 30 min. After washing with PBS, cells were incubated with the primary anti-SERPINB13 antibody (HPA057129, Merck KGaA, Darmstadt, Germany) diluted 1:300 in 0.1 % BSA/PBS for 1 h at RT. After repeated washing with PBS, cells were incubated with the secondary Alexa Fluor®488 antibody (A-11008, Invitrogen, Thermo Fisher Scientific, Waltham, MA, USA) and simultaneous nuclear counterstain was performed for 30 min with Hoechst 33342 (1:1000 in 0.1 % BSA/PBS). Cells were washed again with PBS and subjected to an additional fixation step with 4 % paraformaldehyde for 5 min. After a final wash with PBS, cells were stored in ibidi Mounting Medium until image acquisition on the confocal microscope TCS SP5 (Leica, Wetzlar, Germany).

### RT-qPCR

Total RNA was extracted from LUDLU-1 using innuPREP RNA Mini Kit 2.0 (Analytik Jena, Jena, Germany) according to manufactureŕs protocol. RNA was reverse transcribed into cDNA using Superscript III Reverse Transcriptase (Thermo Fisher Scientific). Real-time PCR was performed on LightCycler 2.0 with the LightCycler 480 SYBR Green I Master (Roche, Basel, Switzerland) completed with specific primers, accordingly: GCTTCTGCCCAACGACATCG SERPINB13-for, ACCGTCCTCCACCTCAAACC SERPINB13-rev, GCTGGCCCATAGTGATCTTT TBP-for, TCCTTGGGTTATCTTCACACG TBP-rev (Thermo Fisher Scientific). Relative quantification was calculated using the software LightCycler® 480 SW 1.5.1 (Roche).

### Statistical Analysis

For methylome and transcriptome analysis, GraphPad Prism 8 (GraphPad Software, Boston,) was used. The results are displayed as mean ±95% confidence interval. The relative gene expression values at the RNA level were quantile-normalized and analyzed using an unpaired t-test and the multiple comparison correction of Benjamini, Krieger, and Yekutieli (Q=5%).

Differential SERPINB13 expression for transcriptome data was calculated in GraphPad Prism 10 with one-way ANOVA and multiple comparison correction of Sidak. Statistics for SERPINB13 gene expression and Western Blot quantification data from *in vitro* experiments were calculated using two-way ANOVA and multiple comparison correction of Sidak in GraphPad Prism 10.

For IHC analysis and data visualization, R software (version 4.0.2, R Foundation, Vienna, Austria; http://www.R-project.org) was used. For survival analysis, the study population was stratified in two (low-high) and four groups (negative, low, moderate, high), respectively, according to Serpin B13 staining intensity assessed conventional-morphologically. The survival was limited to the 5-year overall survival. Survival curves were depicted using Kaplan–Meier plots, and statistical differences were assessed using the log-rank test. To evaluate clinico-pathological parameters as well as immune cell infiltration (hot-cold) and SERPINB13 staining intensity, the Chi-square test was used for categorical data, if the expected cell count was ≥5, else the Fisher’s Exact test was used. The Wilcoxon-rank sum test was used for continuous data. Correlation between SERPINB13 expression and immune cell subtypes was assessed using Spearman correlation. All statistical tests were two-sided, and p-values < 0.05 were considered statistically significant.

We performed Kaplan-Meier analyses in a separate collective of patients (n=780) from data generated using Affymetrix chips and retrieved from kmplot.com. The KM-plotter is an online service capable of assessing the impact of a large number of genes on survival in different cancer types ^18^. Sources of this data collection include GEO, EGA, and TCGA.

## Results

### SERPINB13 is epigenetically modified in NSCLC

Methylome analysis comprised 32 NSCLC samples, including 18 LUSC (14 males, 4 females) and 14 LUAD (5 males, 9 females). Concerning methylation of SERPINB13 CpG loci, we noticed a clear difference between the two NSCLC subtypes as well as tumor-free lung tissues (Figure 1). In particular, LUSC showed a remarkable hypomethylation in 6 of the 8 CpG loci (the top six in hierarchical clustering), while the remaining 2 CpG loci (the bottom two) showed a slight increase in methylation levels. In LUAD, results were not as conclusive. While CpG hypomethylation was observed in some LUAD cases, other cases showed no significant changes in CpG methylation compared to the control group.

**Figure 1.**
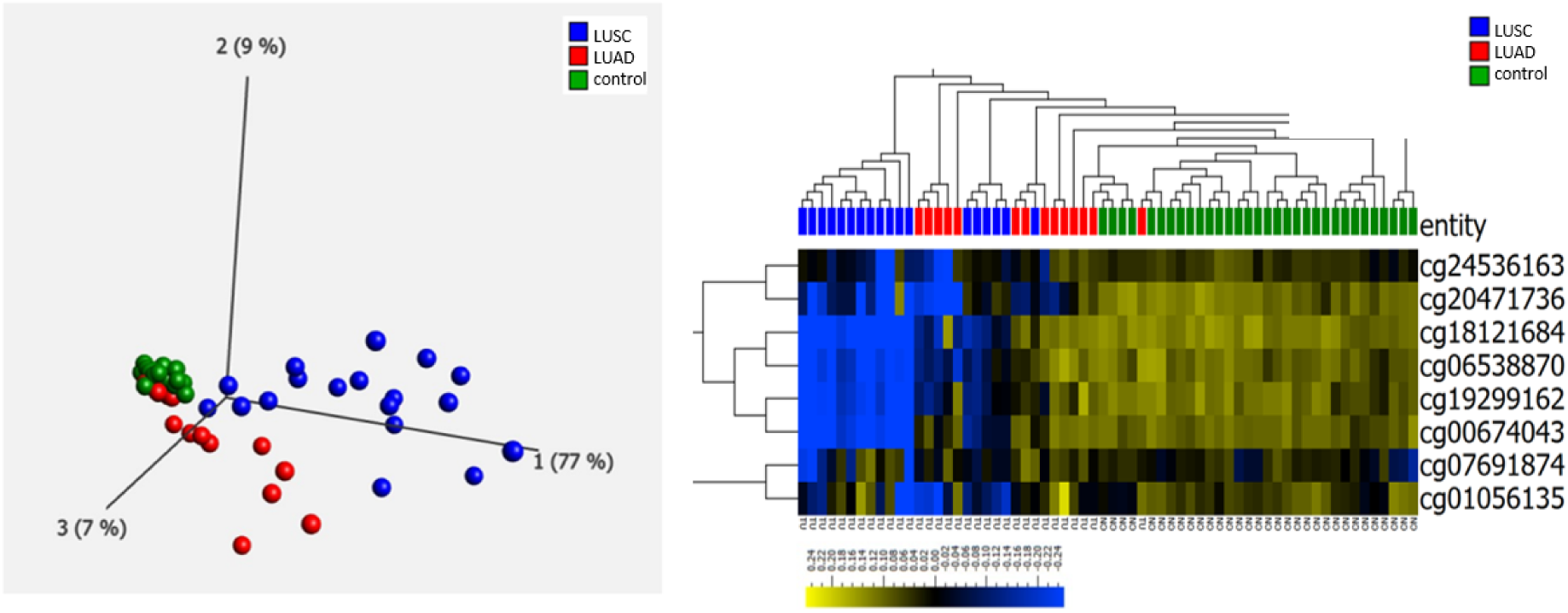
Left: Principal component analysis (PCA) of the methylation profiles from 18 LUSC cases (blue dots), 14 LUAD cases (red dots) and corresponding tumor-free lung tissue (green dots) (n=32; mean DNA methylation = 0). Right: Hierarchical clustering of the methylation levels of SERPINB13 CpG loci using human Methylation 450k BeadChip (upper line: green, tumor-free lung tissue; red, LUAD, blue, LUSC; heatmap: yellow-green, high; blue, low DNA methylation values; mean DNA methylation = 0; range: -0.25-0.25). The target IDs of the CpG loci and their localization (chr.19q) are named in the right-hand row.

### RNA expression of SERPINB13 is upregulated in LUSC

The epigenetic alterations observed in *SERPINB13* CpG loci in NSCLC suggested a potential functional impact on transcriptional regulation. To adress this, we performed targeted RNA expression analysis of tumor and matched tumor-free lung tissue samples from the same patients included in the methylome cohort (n = 18 comprised 10 LUAD and 8 lung LUSC cases). Statistical analysis revealed a strong upregulation of SERPINB13 mRNA expression in LUSC tumors compared to matched tumor-free lung tissues (p ≤ 0.0001). In contrast, *SERPINB13* expression was significantly downregulated in LUAD relative to matched adjacent non-tumor tissues (p ≤ 0.005). Notably, no significant difference in *SERPINB13* expression levels was observed between tumor-free lung tissues from LUSC and LUAD patients (Figure 2).

**Figure 2.**
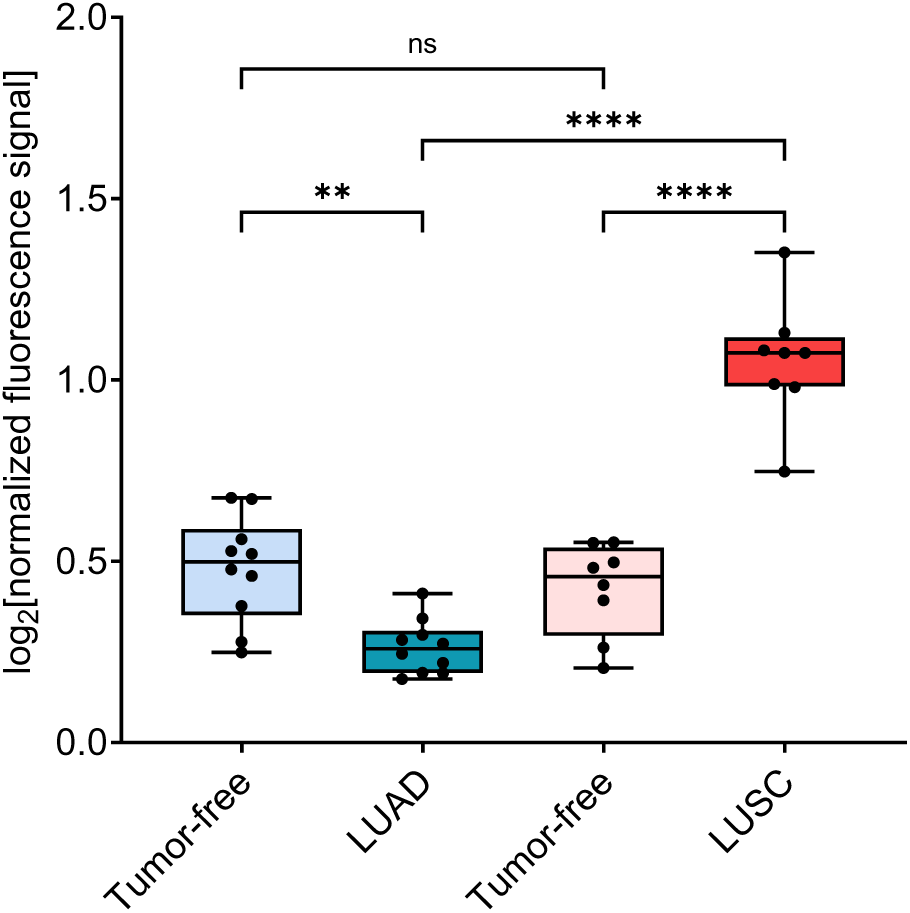
SERPINB13 expression levels were assessed using microarray analysis and visualized as box plots (minimum to maximum values). Fluorescence intensities from tumor-free lung tissues and matched tumor samples (lung adenocarcinoma (LUAD; n=10), lung squamous cell carcinoma (LUSC; n=8)) were percentile-normalized and log₂-transformed. Statistical significance was evaluated using one-way ANOVA ((p ≤ 0.0001, ****), (p ≤ 0.005, **), (ns = not significant)).

### SERPINB13 protein expression pattern

We next performed IHC staining on LUAD and LUSC cases in an independent cohort since the divergent methylome- and transcriptome results for *SERPINB13* expression in both NSCLC subtypes indicated a validation at the protein level. In LUAD, no to a negligible protein expression was detectable (Supplement Figure 1), which was consistent with the significantly lower *SERPINB13* gene expression compared to LUSC and tumor-free lung tissues.

**Supplement Figure 1.**
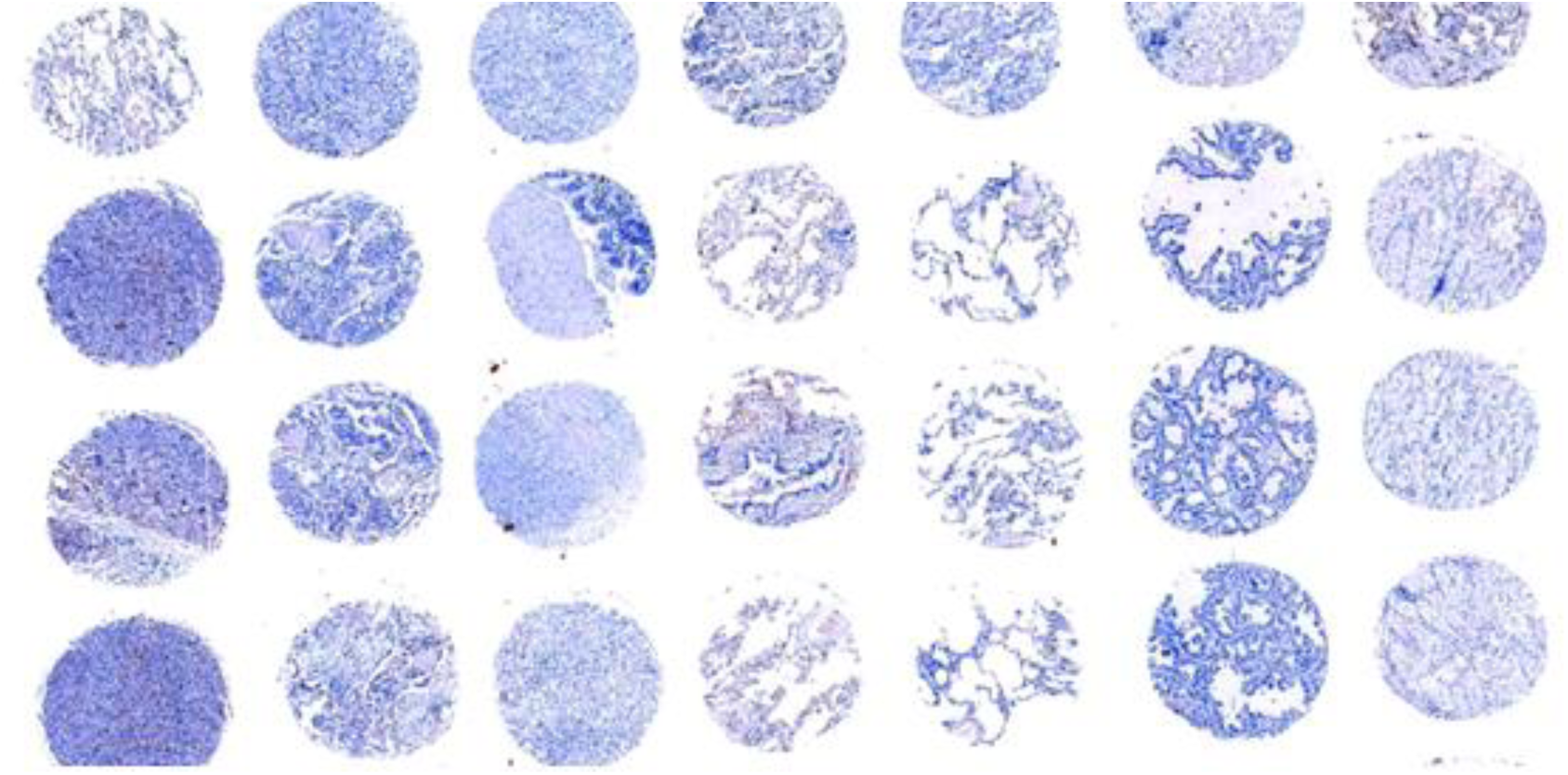
Image of a TMA deriving from LUAD samples showing absent SerpinB13 protein expression

In contrast, a clearly increased protein expression of SERPINB13 was confirmed in LUSC, which is why we focused on this sub-cohort for subsequent analyses concerning correlation analysis of SERPINB13 protein expression with clinico-pathological parameters and immune cell infiltration (Table 1). We observed a homogeneous expression across tissue cores from the same tumor, indicating negligible intratumoral heterogeneity. Across tumor specimens, expression intensity ranged from weak to high levels (62%), besides negative (38%) cases. Overall, 8.7% of LUSC samples showed strong expression (11 out 126), 19 % moderate expression (24 out of 126), 34% weak expression (43 out of 126), and 38.1% (48 out of 126) showed no protein expression of SERPINB13. After dichotomization in a low (negative and weak staining intensity) and high (moderate and strong staining intensity) expressing group, 91 samples (72%) were assigned to the low and 35 samples (28%) to the high expressing group. The distribution of SERPINB13 expression levels differed significantly (p<0.001, Table 1). We stated a variable expression of SERPINB13 in respiratory epithelia with negative cases as well as positive cases demonstrating staining in basal layers. No SERPINB13 expression in pneumocytes was observed. Exemplary IHC images are shown in Figure 3.

**Table 1.** Overview of clinico-pathological characteristics of SERPINB13 high and low expressing group Tumor grading was performed according to the 2015 World Health Organization (WHO) Classification of Lung Tumors^19^. Pathological staging was assigned according to the 8th and 9th edition of the UICC/TNM classification.

| Characteristic | Overall (n=126) | SERPINB13 low (n=91) | SERPINB13 high (n=35) | p-value |
| --- | --- | --- | --- | --- |
| **Sex** |  |  |  | 0.2 |
| female | 43 (34%) | 34 (37%) | 9 (26%) |  |
| male | 83 (66%) | 57 (63%) | 26 (74%) |  |
| **Age** | 68 (61. 73) | 68 (61. 74) | 68 (61. 72) | 0.5 |
| **T-Stage** |  |  |  | 0.8 |
| 1 or 2 | 87 (70%) | 62 (69%) | 25 (71%) |  |
| 3 or 4 | 38 (30%) | 28 (31%) | 10 (29%) |  |
| Missing | 1 | 1 | 0 |  |
| **N-Status** |  |  |  | 0.4 |
| - | 87 (81%) | 58 (78%) | 29 (85%) |  |
| + | 21 (19%) | 16 (22%) | 5 (15%) |  |
| Missing | 18 | 17 | 1 |  |
| **M-Status** |  |  |  | 0.5 |
| - | 44 (94%) | 28 (90%) | 16 (100%) |  |
| + | 3 (6.4%) | 3 (9.7%) | 0 (0%) |  |
| Missing | 79 | 60 | 19 |  |
| **Grading** |  |  |  | 0.2 |
| 2 | 54 (43%) | 42 (47%) | 12 (34%) |  |
| 3 | 71 (57%) | 48 (53%) | 23 (66%) |  |
| Missing | 1 | 1 | 0 |  |
| **UICC 9th Ed.** |  |  |  | 0.13 |
| I or II | 86 (80%) | 56 (76%) | 30 (88%) |  |
| III or IV | 22 (20%) | 18 (24%) | 4 (12%) |  |
| Missing | 18 | 17 | 1 |  |
| **UICC 8th Ed.** |  |  |  | 0.11 |
| I or II | 86 (79%) | 56 (75%) | 30 (88%) |  |
| III or IV | 23 (21%) | 19 (25%) | 4 (12%) |  |
| Missing | 17 | 16 | 1 |  |
| **SerpinB13 (Graduated)** |  |  |  | <0.001 |
| negative | 48 (38%) | 48 (53%) | 0 (0%) |  |
| weak | 43 (34%) | 43 (47%) | 0 (0%) |  |
| moderate | 24 (19%) | 0 (0%) | 24 (69%) |  |
| strong | 11 (8.7%) | 0 (0%) | 11 (31%) |  |

**Figure 3.**
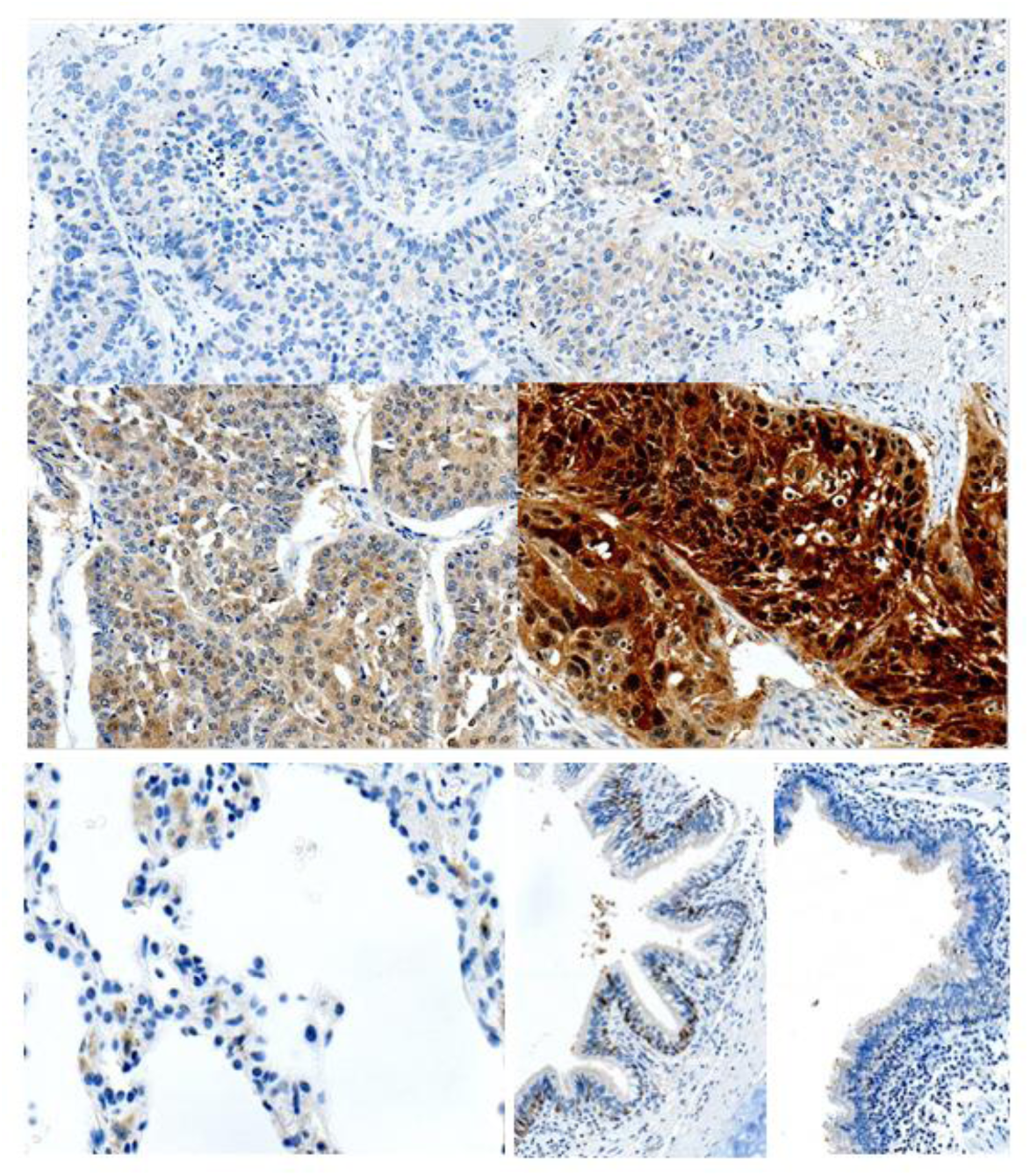
Exemplary images of SERPINB13 protein expression patterns in LUSC. Top left: LUSC with negative expression, top right: LUSC with low expression (both classified as low expressing). Middle left: LUSC with moderate expression, middle right: LUSC with strong expression (both classified as high expressing). The figures demonstrate cytoplasmic staining that appears homogenous within the cores. Bottom left: lung tissue showing no SERPINB13 expression on pneumocytes, bottom middle and right: bronchus epithelium showing variable SERPINB13 expression. (Objective magnification × 400).

### SERPINB13 protein expression in LUSC is positively associated with overall survival

For survival analysis, Kaplan-Meier plots were generated, and log-rank testing was applied (Figure 4). We had access to the relevant survival data from 103 LUSC patients. Samples were first stratified into high and low. Patients with high SERPINB13 protein expression showed significantly better 5-year overall survival than those with low SERPINB13 protein expression (p=0.021), with survival curves diverging early and remaining separated throughout follow-up. After further subdividing expression data into four groups (missing, low, moderate, high expression), results remained statistically significant (p=0.01). The results showed similar favorable survival rates for LUSC with high and moderate SERPINB13 expression, and poor survival rates for cases with low or absent SERPINB13 expression.

**Figure 4.**
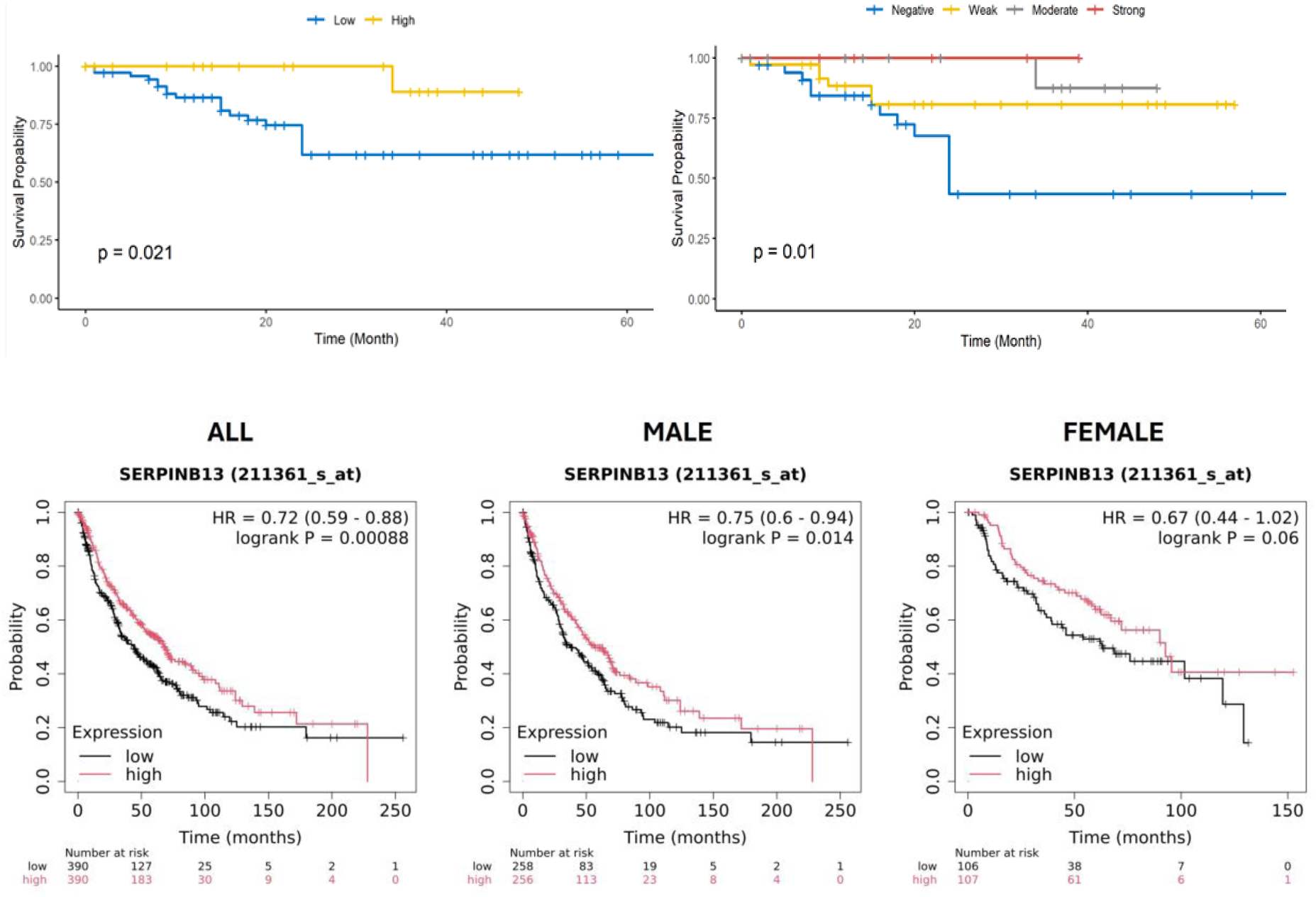
Kaplan-Meier graphs generated from our IHC cohort of LUSC with p-values acquired from log-rank tests after (top left) dichotomization in low and high SERPINB13 expression and (top right bottom) 4-level graduation in negative, weak, moderate and strong Serpin B13 expression. Bottom: Kaplan-Meier survival analysis using mRNA data from LUSC (n=780) retrieved from https://kmplot.com/ analysis/index.php? p=service&cancer=lung, generated using Affymetrix chips (black, low expression; red, high expression). The results are stratified by sex. Statistical evaluation is depicted as a hazard ratio (HR) with 95% confidence intervals and P-value of the log-rank test (log-rank P).

Additionally, we performed Kaplan-Meier analyses in a separate collective of patients (n=780) using kmplot.com (Figure 4). This analysis based on *SERPINB13* gene expression ^18^, yielded in similar results with a significantly better OS for high *SERPINB13* expressing LUSC (p=0.00088).

### Correlation of SERPINB13 protein expression with clinico-pathological characteristics beyond survival

Staining results were correlated with sex, age, T-Stage, N-Stage, M-Stage, UICC-Stage, and tumor grading (Table 1). The distribution of UICC-Stage (T1–2 vs. T3–4) as per 8th and 9th edition did not differ significantly between groups after dichotomization (p=0.11 and 0.13, respectively). Similar results were found for the distribution of T-Stage (p=0.8), N-Stage (p= 0.4) as well as M-Stage (0.5). The grading of tumors and patientś sex and age were also similar between groups (p=0.2, respectively and 0.5).

### SERPINB13 protein expression is linked to immune cell infiltration

During microscopic assessment, varying degrees of immune cell infiltration became obvious. This led us to classify the tumor samples into those with strong immune cell infiltration (“hot”) and those with low immune cell infiltration (“cold”), as previously described ^14^. Figure 5 (left) shows representative images.

**Figure 5.**
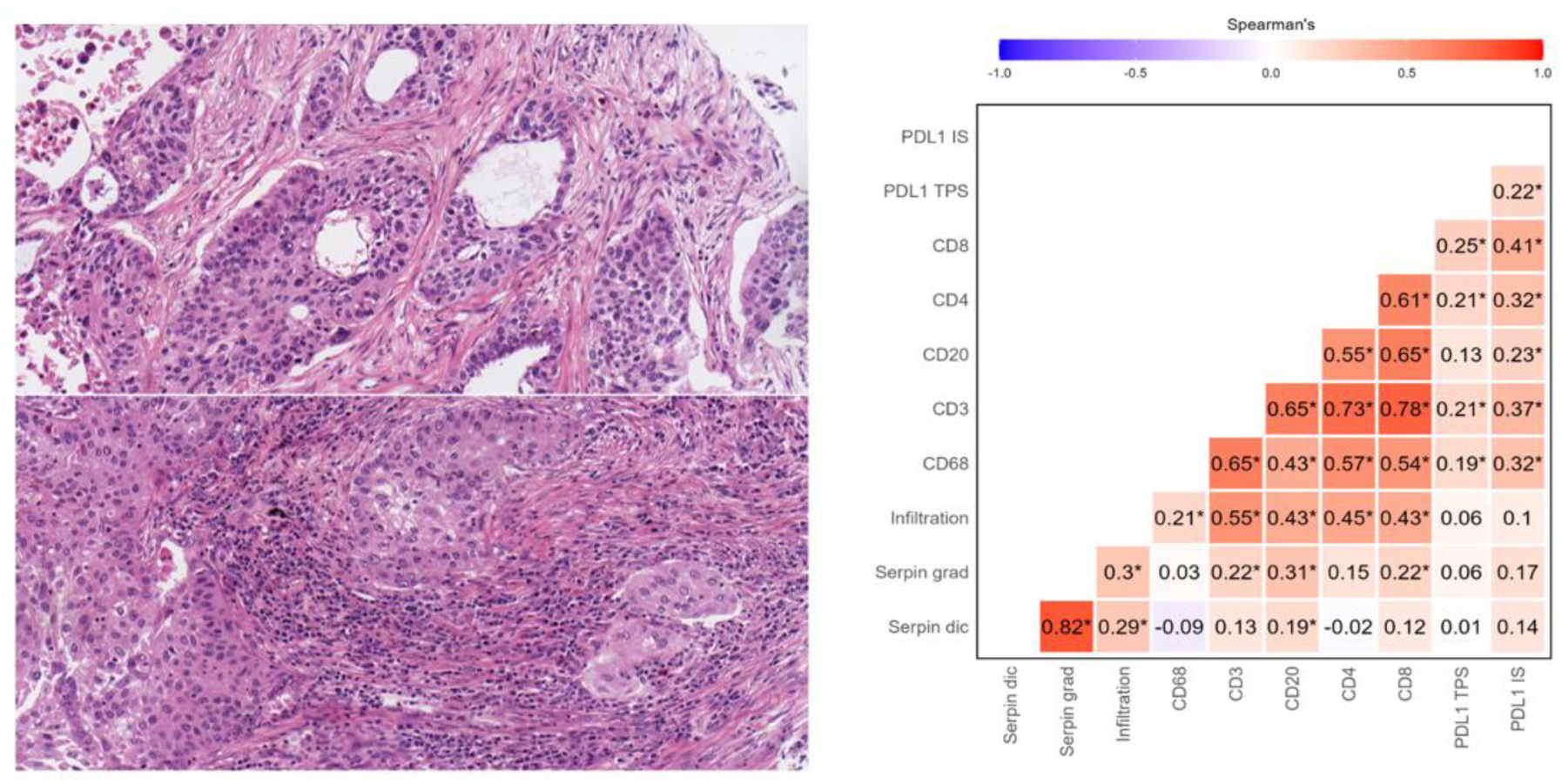
Representative image of a LUSC with low (“cold”) immune cell infiltration (top left) and with strong (“hot”) immune cell infiltration (bottom left) (Objective magnification × 400). Right: Spearman correlation showing the relationship between SERPINB13 protein expression and extent of immune cell subtypes as well as PD-L1-Status meaning tumor proportion score (TPS) and immune score (IS). Significant correlation coefficients are marked with * (p ≤ 0.05)

In our cohort, we classified 77 (61.6%) LUSC as „cold“ and 48 (38.4%) LUSC as „hot“, respectively. When SERPINB13 expression was examined in relation to immune cell infiltration, it became obvious that, within the group of LUSC with dichotomized low SERPINB13 protein expression, there was a proportionally higher number of cases with„cold“ phenotypes (61 out of 90; 67.7%). Moreover, in the group of LUCSs with dichotomized high SERPINB13 protein expression, the proportion of cases with the „hot“ phenotype was higher (19 out of 35; 54.3%). This distribution differed significantly (p=0.023). When subdividing the SERPINB13 expression data as 4-scale (negative, weak, moderate, strong expression), the results remained statistically significant (p=0.013) (Table 2).

**Table 2.** Correlation of SERPINB13 protein expression with immune cell infiltration.

| SERPINB13 expression | cold (n=77) | hot (n=48) | total (n=125)<br>missing:1 | p-value |
| --- | --- | --- | --- | --- |
| **dichotomized** |  |  |  | 0.023 |
| low | 61 (79.2%) | 29 (60.4%) | 90 (72.0%) |  |
| high | 16 (20.8%) | 19 (39.6%) | 35 (28.0%) |  |
| **graduated** |  |  |  | 0.013 |
| negative | 37 (48.1%) | 10 (20.8%) | 47 (37.6%) |  |
| weak | 24 (31.2%) | 19 (39.6%) | 43 (34.4%) |  |
| moderate | 12 (15.6%) | 12 (25.0%) | 24 (19.2%) |  |
| strong | 4 (5.2%) | 7 (14.6%) | 11 (8.8%) |  |

SERPINB13 low expressing cases were associated with a poorer prognosis, whilst high expressing cases showed a favorable prognosis (Figure 4). We also carried out a survival analysis focusing solely on immune cell infiltration, independent of SERPINB13 expression, and here we also found that LUSC with a cold phenotype had a poorer prognosis, whilst LUSC cases with hot phenotype had a more favorable prognosis (p=0.00092; Supplement Figure 2). The results therefore suggest a link between SERPINB13 protein expression and immune cell infiltration as well as overall survival.

**Supplement Figure 2.**
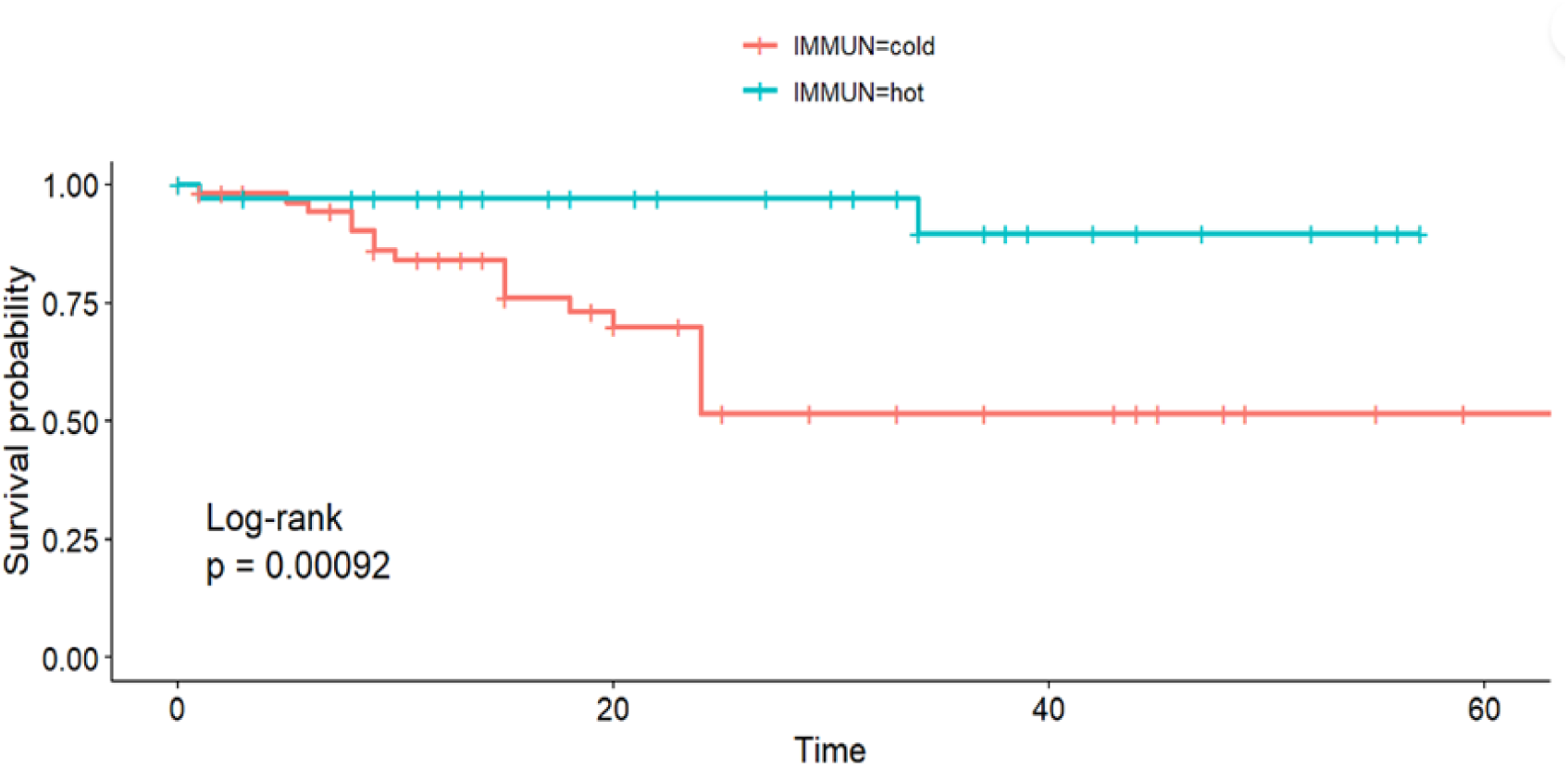
Kaplan-Meier graphs with p-values acquired from log-rank tests of LUSC with low (“cold”) immune cell infiltration and with strong (“hot”) immune cell infiltration.

Besides the pure extent of immune cell infiltration indicated as „hot“ and „cold“, we further correlated SERPINB13 expression and immune cell subtypes including macrophages, CD3-, CD20-, CD8- and CD4-positive lymphocytes as well as PD-L1- and PD-1-Status. We found a positive correlation between the graduated protein expression of SERPINB13 with all parameters, which proved significant for the extent of immune cell infiltration, CD3-positive T-lymphocytes, CD8-positive cytotoxic T-lymphocytes, and CD20-positive B-lymphocytes. In dichotomised SERPINB13 expression data a positive correlation was found for all parameters except for macrophages (CD68), and this was significant solely for the extent of immune cell infiltration and CD20-positive B-lymphocytes (Figure 5; right).

### Transwell co-cultures with PBMCs show a modulation of SERPINB13 expression in LUSC by secreted mediators in a time-dependent manner

Co-culture experiments using the SERPINB13-expressing LUSC cell line LUDLU-1 and different concentrations of peripheral blood mononuclear cells (PBMCs) revealed a time- and concentration-dependent regulation of SERPINB13 expression (Figure 6). After 1 day of co-culture, a significant increase in SERPINB13 expression in LUDLU-1 cells was observed in comparison to the control group without PBMCs after the addition of 500,000 and 200,000 PBMCs. At lower PBMC concentrations (50,000 PBMCs), a slight increase in SERPINB13 expression was also detected; however, this effect did not reach statistical significance. In contrast, after 5 days of incubation, the effect was reversed. Compared to the control group, SERPINB13 expression decreased in a PBMC-dependent manner with increasing PBMC numbers. The strongest effect was observed in co-cultures containing 500,000 PBMCs, where a significant reduction in SERPINB13 expression was detected. These dynamic changes of SERPINB13 were confirmed additionally on protein level by Western Blot analysis (Figure 6). Overall, these findings indicate a dynamic and time-dependent regulation of SERPINB13 expression mediated by interactions with immune cells.

**Figure 6.**
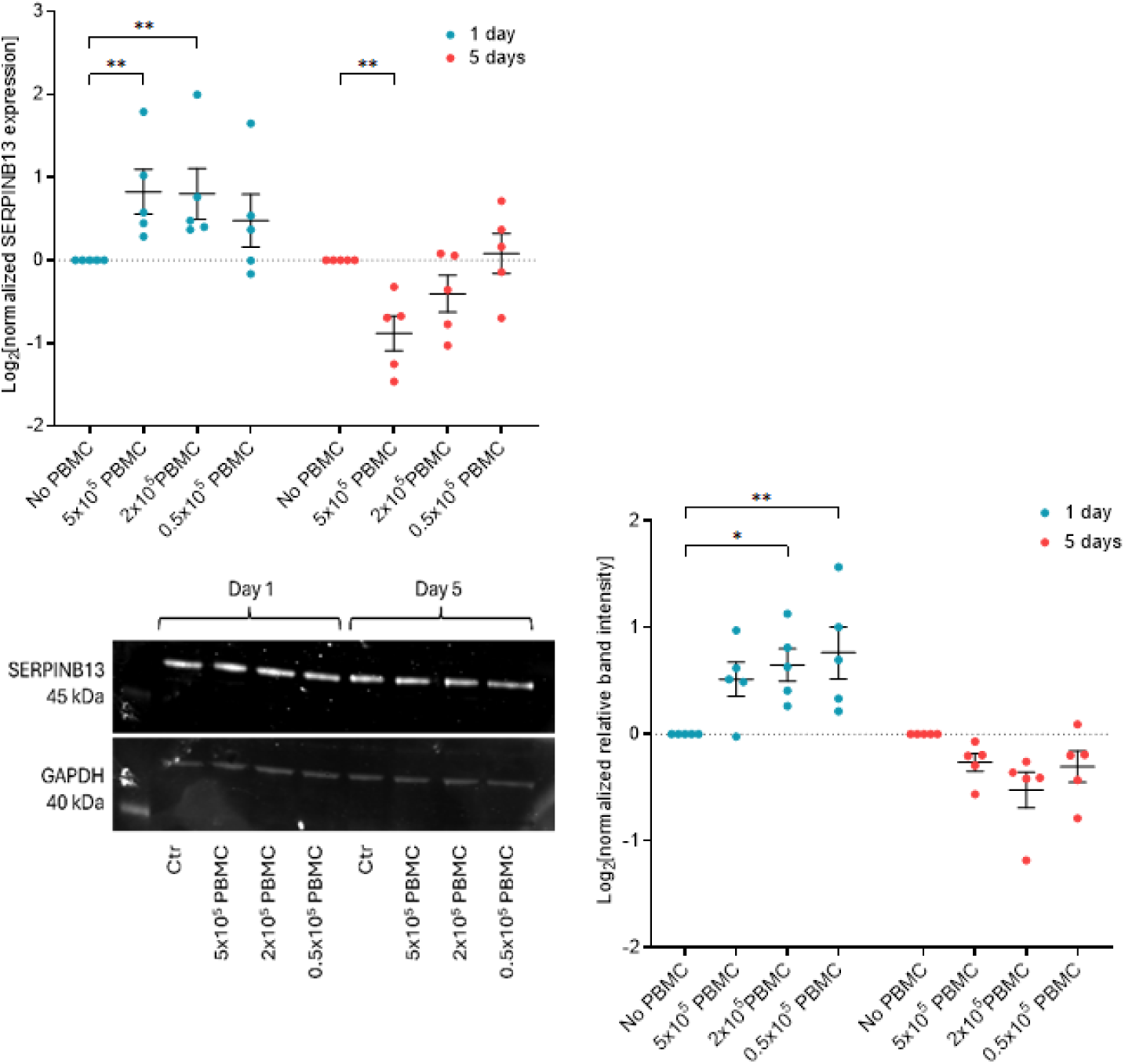
Top: SERPINB13 expression in LUDLU-1 cells co-cultured with PBMCs. LUDLU-1 cells were cultured in transwell inserts with PBMCs from five donors. SERPINB13 mRNA levels were measured by RT-qPCR at 1 day and 5 days post-co-culture, normalized to TBP and relative to controls (no PBMCs). Bottom: SERPINB13 protein expression in LUDLU-1 cells co-cultured with PBMCs. Cells were co-cultured in transwell inserts with PBMCs from five donors for 1 and 5 days. SERPINB13 levels were analyzed by Western blot, with GAPDH as loading control. Representative image is shown (bottom left). Bottom right: Band intensities were estimated and values were normalized to control to quantify amount of SERPINB13 protein. Log₂-transformed values in scatter dot plot graphs are presented as mean ± SEM (n = 5 donors) and statistic was calculated by two-way ANOVA (p ≤ 0.05, *; p ≤ 0.01, **).

### Confocal microscopy shows cytoplasmic subcellular staining pattern of SERPINB13 in LUDLU-1 cells

Immunofluorescence staining was performed to assess the subcellular localization of SERPINB13 in LUDLU-1 cells. SERPINB13-specific signal (green) was detected throughout the cytoplasm, with a predominantly diffuse distribution and punctate accumulations. Notably, SERPINB13 signal was largely excluded from the nucleus (red), indicating a cytoplasmic localization of the protein. No specific signal was observed in the secondary antibody-only control (lower row), confirming the specificity of the staining (Supplement Figure 3).

**Supplement Figure 3.**
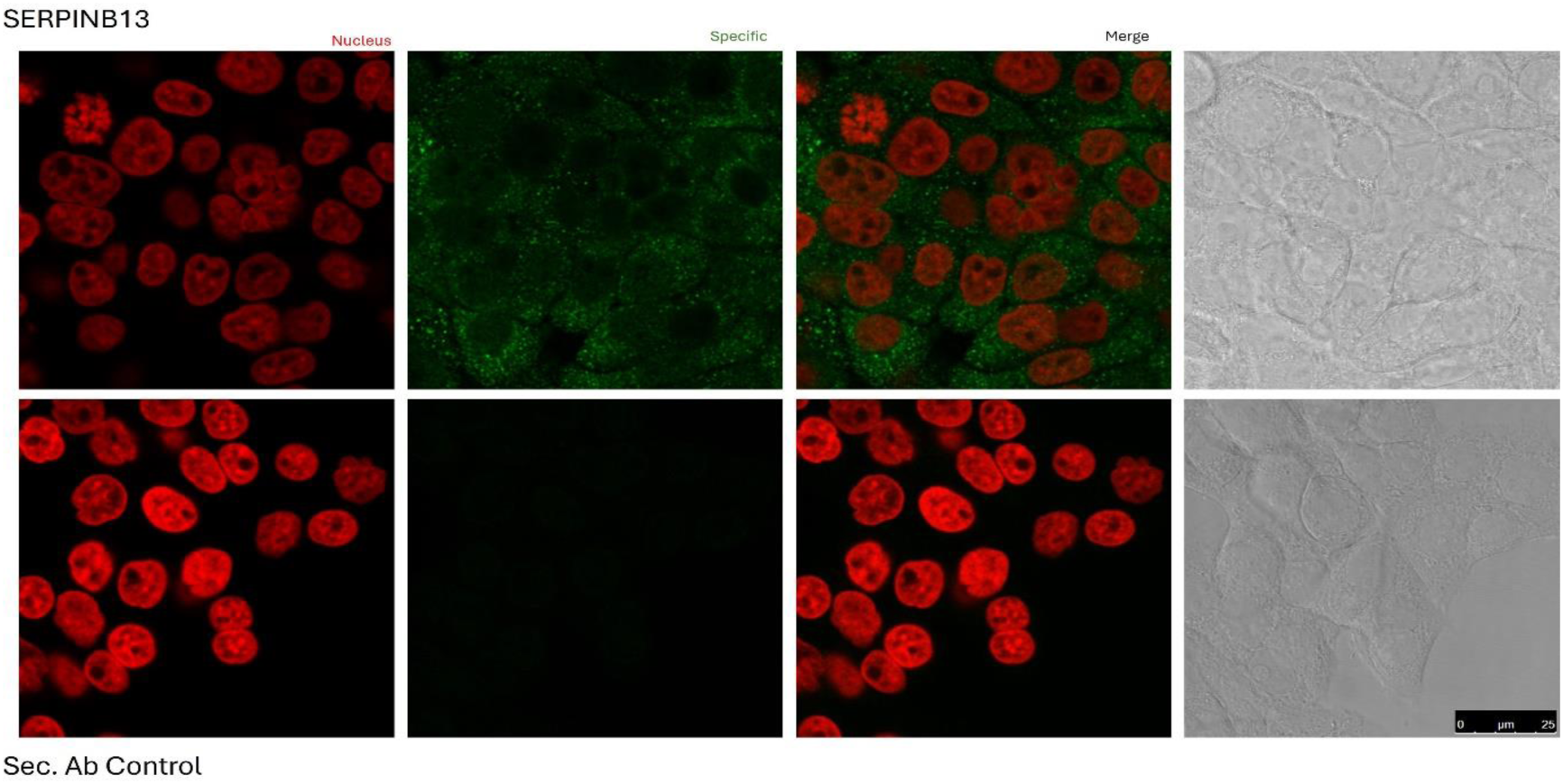
Immunofluorescence staining of SERPINB13 in LUDLU-1 cells. Representative confocal immunofluorescence images of LUDLU-1 cells stained for SERPINB13 (green) and nuclei (red). Upper row: cells incubated with primary anti-SERPINB13 antibody followed secondary antibody. Lower row: secondary antibody-only control. From left to right: nuclear channel, channel for antibody staining, merged image, and phase contrast. Scale bar: 25 µm.

## Discussion

As part of a large scaled study, aimed at illuminating at least some of the complex events taking place in the interaction of NSCLC tumor cells and the tumor microenvironment, we identified the serine protease inhibitor SERPINB13 as a promising candidate gene by transcriptome and methylome analyses. Comparably little was known about SERPINB13 in NSCLC, what prompted us to undertake this investigation.

### SERPINB13 exhibits subtype-specific regulation in LUAD and LUSC

First, we investigated the epigenetic modification of *SERPINB13* CpG loci in NSCLC compared to tumor-free lung tissues. We identified a clear difference between the two NSCLC subtypes, LUAD and LUSC. Specifically, in LUSC, we observed a remarkable hypomethylation of several analyzed CpG loci.

The assessed epigenetic alterations at SERPINB13 CpG loci in NSCLC suggested a potential functional impact on transcriptional regulation. For this reason, we continued with transcriptome analyses on NSCLC cases in the same cohort. Here, we could state a significant upregulation of SERPINB13 mRNA expression in LUSC cases compared to corresponding normal lung tissue, while a significant downregulation was observed in LUAD. Since we found no significant difference in SERPINB13 expression levels between tumor-free lung tissues from LUSC and LUAD samples, it can be concluded that the differential expression pattern is specific to the tumor subtype. This observation provides strong evidence for subtype-specific transcriptional regulation of SERPINB13 in NSCLC, likely driven by epigenetic mechanisms such as the observed promoter hypomethylation in LUSC, suggesting that *SERPINB13* expression is linked to subtype-specific programs in NSCLC (Figure 1 and 2).

### SERPINB13 is associated with favorable survival

To validate the transcriptome findings, we assessed SERPINB13 protein expression using IHC on a larger and independent cohort of NSCLC. In line with the transcriptome data, no significant SERPINB13 protein expression was detected in LUAD. In contrast, and corroborating the RNA results, we confirmed protein expression of SERPINB13 in LUSC, ranging from weak to high levels in most cases. Survival analysis revealed a significantly better 5-year OS for LUSC patients with high SERPINB13 protein expression compared to those with low expression (Figure 4). This was also the case when protein expression was stratified into four categories (negative, weak, moderate, and high). Since we did not identify any significant correlation with other clinico-pathological data, SERPINB13 seems to be expressed independently of these.

This is the first study to report on SERPINB13 protein expression in NSCLC. Therefore, direct comparison with existing literature is limited. However, studies have investigated SERPINB13 expression in SCC of head and neck (HNSCC) and oral cavity (OSCC) ^7,8^. Given that SERPINB13 is primarily expressed on keratinocytes and plays a role in squamous cell carcinomas, a comparison is relevant despite their different anatomical locations. Furthermore, LUSC, HNSCC, and OSCC share common risk factors, notably smoking ^20^. A study on HNSCC, similar in size to our cohort, reported that SERPINB13 protein expression was downregulated in most cases compared to non-neoplastic epithelium, and this downregulation was significantly associated with decreased DFS and OS. These results are comparable to ours, as we categorized most LUSC cases as low expressing (Table 1), and low expression was associated with adverse OS (Figure 4). Due to insufficient follow-up data in our cohort, an analysis correlating SERPINB13 expression with DFS could not be performed. Another study by Shellenberger et al. examined SERPINB13 protein expression in a cohort of SCC from various upper aerodigestive tract sites and found weak SERPNB13 protein expression in the vast majority ^4^.

To address the lack of data on correlation of SERPINB13 protein expression and survival in NSCLC cohorts in literature, we validated our survival results with publicly available mRNA data from another independent cohort obtained from kmplot.com. Notably, results were similar with our dichotomized analysis, showing a favorable prognosis for LUSC patients with high *SERPINB13* mRNA expression (Figure 4),consistent with a study from Relli et al. ^9^.

A study on OSCC did not include any SERPINB13 protein expression data either but found significant downregulation of *SERPINB13* mRNA, interpreted as a sign for a tumor suppressive role. However, the reference tissue used was different (squamous epithelium vs. lung tissue), making the data not directly comparable^8^. The study by Zhang et al. reported mRNA expression results in line with ours, showing SERPINB13 expression at higher levels in LUSC tissues compared to adjacent non-cancerous samples ^12^. Both results might indicate that *SERPINB13* harbors a specific biological relevance for LUSC.

One aspect is additionally worth mentioning. SERPINB13 protein expression was reported on respiratory epithelium of the trachea and bronchus, besides other tissues, without specifying the number of cases ^7^. Here, the authors stated weaker expression levels compared to that seen in the squamous epithelia. We stated a variable expression of SERPINB13 in basal layers of respiratory epithelium (Figure 3). Given that LUSC derived from squamous dysplasia, which arises from squamous metaplasia of respiratory epithelium, it could be assumed that there is an etiological link between SERPINB13 expression in respiratory stem cells and squamous differentiation. Notably, we found no SERPINB13 protein expression in pneumocytes. This might also be the reason why we did not find any SERPINB13 expression in the LUAD subcohort.

### SERPINB13 expression is linked to an immune-inflamed tumor microenvironment in LUSC

Given the involvement of SERPINB13 in inflammatory skin diseases^6^ and the varying degree of immune cell infiltration in NSCLC, we analyzed its protein expression in relation to tumor associated immune cells. We found the proportion of cases with the „hot“ phenotype to be higher within the group of LUSC with high SERPINB13 protein expression, and vice versa. This distribution differed significantly (Figure 5, Table 2). These findings are intriguing, considering that SERPINB13 expression correlated with OS, with the high SERPINB13-expressing LUSC group, characterised by a predominance of hot TIME phenotype, also represented the group with better OS, and vice versa. Thus, our data suggest a link between tumor SERPINB13 expression, TIME, and prognosis of LUSC.

It is well established that TIME phenotype alone has prognostic value^2^ , and we confirmed this for our IHC cohort as well (Supplement Figure 2). Nevertheless, *in silico* data published by Zhang et al. revealed a negative correlation between SERPINB13 RNA-expression and the immune score but a positive correlation of SERPINB13 expression and improved prognosis in LUSC ^12^. These results are in part in line with ours as we also stated a positive correlation between SERPINB13 expression and survival, but additionally with tumor associated immune cell infiltration. This can be explained since we addressed the protein level, (Table 2, Figure 4, Supplement Figure 2).

We further correlated SERPINB13 protein expression and several immune cell subtypes, as well as PD-L1-Status. Here, we found a positive correlation between dichotomized SERPINB13 protein expression and all parameters except for macrophages and CD4-positive T-lymphocytes, which proved significant for CD20-positive B-lymphocytes, and found minor variations for graduated SERPINB13 expression (Figure 5). This is in line with the co-culture experiments applying PBMCs and might indicate that tumor infiltrating macrophages do not drive SERPINB13.

### Dynamic regulation of SERPINB13 by immune cells

An intriguing finding of the present study is the association between high SERPINB13 expression and increased immune cell infiltration in patient samples, which was further supported by the PBMC co-culture experiments. To our knowledge, this is the first study demonstrating that immune cells directly modulate SERPINB13 expression in LUSC cells.

Our findings suggest a bidirectional interaction between tumor cells and the immune microenvironment. It is conceivable that an immunologically active (“hot”) tumor microenvironment promotes SERPINB13 expression through inflammatory signaling, thereby linking high SERPINB13 expression to favorable prognosis. Conversely, SERPINB13 may also influence the tumor microenvironment through its inhibitory effects on cathepsin L, a cysteine protease involved in extracellular matrix degradation, tumor invasion, metastasis, and angiogenesis^21^. Inhibition of cathepsin L may therefore represent one mechanism underlying the favorable prognostic impact of SERPINB13 observed in our cohort.

Similar tumor-modulating functions have been described for other members of the SERPIN family. SERPINB5, for example, has been associated with reduced invasion, decreased metastatic potential, and improved clinical outcome in several epithelial malignancies ^22, 23^. Likewise, other clade B SERPINs, for example SERPINB2 and SERPINB3/B4, are regulated by inflammatory stimuli and participate in shaping the interaction between tumor cells and the immune microenvironment^24,25,26^. Although the biological functions of these SERPINs differ, these observations suggest that immune responsiveness may represent a common feature of intracellular SERPIN family members. Our findings extend this concept to SERPINB13 and support a previously unrecognized role in the regulation of the LUSC tumor immune microenvironment.

The significant induction of SERPINB13 after one day of PBMC co-culture suggests that immune-derived signals stimulate SERPINB13 expression as part of an early adaptive response. Since SERPINB13 has been reported to protect epithelial cells from cathepsin L-mediated apoptosis, this transient upregulation may represent a cytoprotective response to immune-mediated stress ^5^. In contrast, prolonged co-culture resulted in reduced SERPINB13 expression, which may reflect diminished immune cell activity under in vitro conditions, tumor cell adaptation, or changes in the balance between survival and cell death pathways. However, whether SERPINB13 directly regulates apoptosis in LUSC cells remains unknown.

Further mechanistic studies are needed to identify the immune cell populations and signaling pathways regulating SERPINB13 expression and to determine whether this interaction contributes to the response to immune checkpoint inhibition in lung squamous cell carcinoma.

## Limitations

Our study has several limitations. Firstly, our methylome/transcriptome and protein data do not derive from the same cohort, making direct comparisons challenging. The size of our LUSC subcohort limits our ability to draw definitive conclusions regarding prognosis. Another limitation is the age of the cohort, as immune checkpoint inhibitor (ICI) therapy was not routinely administered to these patients, preventing us from correlating our SERPINB13 expression results with ICI treatment response. While our findings suggest an association, they do not establish a causal role of SERPINB13 expression and improved survival. We further demonstrated an association between SERPINB13 expression and immune cells in vitro. However, the in vitro co-culture system only partially reflects the complexity of the tumor immune microenvironment and the mechanisms driving this interaction remain unknown.

**In summary**, SERPINB13 is a promising prognostic biomarker for LUSC and is linked to an immune-inflamed tumor microenvironment. This could have potential implications for the development of immunomodulatory strategies and personalized therapeutic approaches. Given its reliable immunohistochemical detectability and easy evaluation, its incorporation into routine diagnostic appears feasible. Prospective studies with independent validation cohorts are required to confirm results presented in this study.

## Acknowledgements

The authors thank Eva Dreyer, Jasmin Tiebach and Christian Rosero for their excellent technical assistance as well as the BioMaterialBank North and the blood donation services of the Research Center Borstel.

## Ethics approval and consent to participate

The study was conducted in accordance with the Declaration of Helsinki, and the protocol was approved by the local ethics council at the University of Luebeck (AZ-12-220, AZ 16-277). Informed consent was obtained from subjects involved in the study as required by the Ethics Committee.

## Availability of data and materials

The transcriptome and methylome data presented in this study are openly available from the GEO database (GEO accession number GSE74706 for the transcriptome data and GSE75008 for the methylome data). The survival data from a different cohort derived from the TCGA are publicly available and can be found here: kmplot.com. Further data presented in this study are available upon request from the corresponding author. The data is not publicly available due to ethical restrictions.

## Abbreviations

DFS: disease-free survival
FFPE: formalin-fixed paraffin-embedded
HNSCC: head and neck squamous cell carcinoma
IHC: immunohistochemistry
ICI: immune checkpoint inhibitor
IS: immune score
LOH: loss of heterogeneity
LMS: LUSC multi-omics signature
LUAD: lung adenocarcinoma
LUSC: lung squamous cell carcinoma
NSCLC: non small cell lung cancer
OS: overall survival
OSCC: oral squamous cell carcinoma
PBMC: peripheral blood mononuclear cells
PFS: progression free survival
RT-qPCR: quantitative real-time PCR
SCC: squamous cell carcinoma
SERPIN: Serine protease inhibitor
TCGA: The Cancer Genome Atlas
TIME: tumor immune microenvironment
TMA: tissue microarray
TPS: tumor proportion score
UICC: Union for International Cancer Control

## Literature

1. Howlader, N., Forjaz, G., Mooradian, M.J., Meza, R., Kong, C.Y., Cronin, K.A., Mariotto, A.B., Lowy, D.R., and Feuer, E.J. (2020). The Effect of Advances in Lung-Cancer Treatment on Population Mortality. N Engl J Med 383, 640–649. 10.1056/NEJMoa1916623.

2. Yan, Q., Li, S., He, L., and Chen, N. (2024). Prognostic implications of tumor-infiltrating lymphocytes in non-small cell lung cancer: a systematic review and meta-analysis. Front Immunol 15, 1476365. 10.3389/fimmu.2024.1476365.

3. Marwitz, S., Brunnström, H., Gulyas, M., Botling, J., Reck, M., Kugler, C., Elfving, H., Micke, P., Strell, C., and Goldmann, T. (2026). Left behind but not left alone: Excluded cell populations in the non-small cell lung cancer stroma predict superior long-term overall survival. Eur J Cancer 244, 116868. 10.1016/j.ejca.2026.116868.

4. Shellenberger, T.D., Mazumdar, A., Henderson, Y., Briggs, K., Wang, M., Chattopadhyay, C., Jayakumar, A., Frederick, M., and Clayman, G.L. (2005). Headpin: a serpin with endogenous and exogenous suppression of angiogenesis. Cancer Res 65, 11501–11509. 10.1158/0008-5472.CAN-05-2262.

5. Welss, T., Sun, J., Irving, J.A., Blum, R., Smith, A.I., Whisstock, J.C., Pike, R.N., von Mikecz, A., Ruzicka, T., Bird, P.I., et al. (2003). Hurpin is a selective inhibitor of lysosomal cathepsin L and protects keratinocytes from ultraviolet-induced apoptosis. Biochemistry 42, 7381–7389. 10.1021/bi027307q.

6. Moussali, H., Bylaite, M., Welss, T., Abts, H.F., Ruzicka, T., and Walz, M. (2005). Expression of hurpin, a serine proteinase inhibitor, in normal and pathological skin: overexpression and redistribution in psoriasis and cutaneous carcinomas. Exp Dermatol 14, 420–428. 10.1111/j.0906-6705.2005.00300.x.

7. de Koning, P.J.A., Bovenschen, N., Leusink, F.K.J., Broekhuizen, R., Quadir, R., van Gemert, J.T.M., Hordijk, G.J., Chang, W.-S.W., van der Tweel, I., Tilanus, M.G.J., et al. (2009). Downregulation of SERPINB13 expression in head and neck squamous cell carcinomas associates with poor clinical outcome. Int J Cancer 125, 1542–1550. 10.1002/ijc.24507.

8. Shiiba, M., Nomura, H., Shinozuka, K., Saito, K., Kouzu, Y., Kasamatsu, A., Sakamoto, Y., Murano, A., Ono, K., Ogawara, K., et al. (2010). Down-regulated expression of SERPIN genes located on chromosome 18q21 in oral squamous cell carcinomas. Oncol Rep 24, 241–249. 10.3892/or_00000852.

9. Relli, V., Trerotola, M., Guerra, E., and Alberti, S. (2018). Distinct lung cancer subtypes associate to distinct drivers of tumor progression. Oncotarget 9, 35528–35540. 10.18632/oncotarget.26217.

10. Yuan, F., Lu, L., and Zou, Q. (2020). Analysis of gene expression profiles of lung cancer subtypes with machine learning algorithms. Biochim Biophys Acta Mol Basis Dis 1866, 165822. 10.1016/j.bbadis.2020.165822.

11. Zhan, C., Yan, L., Wang, L., Sun, Y., Wang, X., Lin, Z., Zhang, Y., Shi, Y., Jiang, W., and Wang, Q. (2015). Identification of immunohistochemical markers for distinguishing lung adenocarcinoma from squamous cell carcinoma. J Thorac Dis 7, 1398–1405. 10.3978/j.issn.2072-1439.2015.07.25.

12. Zhang, X., Zhang, P., Ren, Q., Li, J., Lin, H., Huang, Y., and Wang, W. (2025). Integrative multi-omic and machine learning approach for prognostic stratification and therapeutic targeting in lung squamous cell carcinoma. Biofactors 51, e2128. 10.1002/biof.2128.

13. Marwitz, S., Kolarova, J., Reck, M., Reinmuth, N., Kugler, C., Schädlich, I., Haake, A., Zabel, P., Vollmer, E., Siebert, R., et al. (2014). The tissue is the issue: improved methylome analysis from paraffin-embedded tissues by application of the HOPE technique. Lab Invest 94, 927–933. 10.1038/labinvest.2014.79.

14. Kirfel, J., Kümpers, C.C., Fähnrich, A., Heidel, C., Jokic, M., Vlasic, I., Marwitz, S., Goldmann, T., Pasternack, H., Bohnet, S., et al. (2021). PD-L1 Dependent Immunogenic Landscape in Hot Lung Adenocarcinomas Identified by Transcriptome Analysis. Cancers (Basel) 13, 4562. 10.3390/cancers13184562.

15. Salcher, S., Sturm, G., Horvath, L., Untergasser, G., Kuempers, C., Fotakis, G., Panizzolo, E., Martowicz, A., Trebo, M., Pall, G., et al. (2022). High-resolution single-cell atlas reveals diversity and plasticity of tissue-resident neutrophils in non-small cell lung cancer. Cancer Cell 40, 1503–1520.e8. 10.1016/j.ccell.2022.10.008.

16. Scheel, A.H., Dietel, M., Heukamp, L.C., Jöhrens, K., Kirchner, T., Reu, S., Rüschoff, J., Schildhaus, H.U., Schirmacher, P., Tiemann, M., et al. (2016). [Predictive PD-L1 immunohistochemistry for non-small cell lung cancer : Current state of the art and experiences of the first German harmonization study]. Pathologe 37, 557–567. 10.1007/s00292-016-0189-1.

17. Boyum, A. (1964). SEPARATION OF WHITE BLOOD CELLS. Nature 204, 793–794. 10.1038/204793a0.

18. Győrffy, B. (2024). Transcriptome-level discovery of survival-associated biomarkers and therapy targets in non-small-cell lung cancer. Br J Pharmacol 181, 362–374. 10.1111/bph.16257.

19. Travis, W.D., Brambilla, E., Nicholson, A.G., Yatabe, Y., Austin, J.H.M., Beasley, M.B., Chirieac, L.R., Dacic, S., Duhig, E., Flieder, D.B., et al. (2015). The 2015 World Health Organization Classification of Lung Tumors: Impact of Genetic, Clinical and Radiologic Advances Since the 2004 Classification. J Thorac Oncol 10, 1243–1260. 10.1097/JTO.0000000000000630.

20. Polo, V., Pasello, G., Frega, S., Favaretto, A., Koussis, H., Conte, P., and Bonanno, L. (2016). Squamous cell carcinomas of the lung and of the head and neck: new insights on molecular characterization. Oncotarget 7, 25050–25063. 10.18632/oncotarget.7732.

21. Fujii, Y., Asadi, Z., and Mehla, K. (2025). Cathepsins: Emerging targets in the tumor ecosystem to overcome cancers. Semin Cancer Biol 112, 150–166. 10.1016/j.semcancer.2025.04.001.

22. He, X., Ma, Y., Huang, Z., Wang, G., Wang, W., Zhang, R., Guo, G., Zhang, X., Wen, Y., and Zhang, L. (2023). SERPINB5 is a prognostic biomarker and promotes proliferation, metastasis and epithelial-mesenchymal transition (EMT) in lung adenocarcinoma. Thorac Cancer 14, 2275–2287. 10.1111/1759-7714.15013.

23. Teoh, S.S.Y., Whisstock, J.C., and Bird, P.I. (2010). Maspin (SERPINB5) is an obligate intracellular serpin. J Biol Chem 285, 10862–10869. 10.1074/jbc.M109.073171.

24. Sun, Y., Sheshadri, N., and Zong, W.-X. (2017). SERPINB3 and B4: From biochemistry to biology. Semin Cell Dev Biol 62, 170–177. 10.1016/j.semcdb.2016.09.005.

25. Catanzaro, J.M., Sheshadri, N., Pan, J.-A., Sun, Y., Shi, C., Li, J., Powers, R.S., Crawford, H.C., and Zong, W.-X. (2014). Oncogenic Ras induces inflammatory cytokine production by upregulating the squamous cell carcinoma antigens SerpinB3/B4. Nat Commun 5, 3729. 10.1038/ncomms4729.

26. Schroder, W.A., Major, L., and Suhrbier, A. (2011). The role of SerpinB2 in immunity. Crit Rev Immunol 31, 15–30. 10.1615/critrevimmunol.v31.i1.20.

